# Slit/Robo signals prevent spinal motor neuron emigration by organizing the spinal cord basement membrane

**DOI:** 10.1101/690859

**Authors:** Minkyung Kim, Clare H Lee, Sarah J Barnum, Roland CJ Watson, Jennifer Li, Grant S Mastick

## Abstract

The developing spinal cord builds a boundary between the CNS and the periphery, in the form of a basement membrane. The spinal cord basement membrane is a barrier that retains CNS neuron cell bodies, while being selectively permeable to specific axon types. Spinal motor neuron cell bodies are located in the ventral neural tube next to the floor plate and project their axons out through the basement membrane to peripheral targets. However, little is known about how spinal motor neuron cell bodies are retained inside the ventral neural tube, while their axons can exit. In previous work, we found that disruption of Slit/Robo signals caused motor neuron emigration outside the spinal cord. In the current study, we investigate how Slit/Robo signals are necessary to keep spinal motor neurons within the neural tube. Our findings show that when Slit/Robo signals were removed from motor neurons, they migrated outside the spinal cord. Furthermore, this emigration was associated with abnormal basement membrane protein expression in the ventral spinal cord. Using Robo2 and Slit2 conditional mutants, we found that motor neuron-derived Slit/Robo signals were required to set up a normal basement membrane in the spinal cord. Together, our results suggest that motor neurons produce Slit signals that are required for the basement membrane assembly to retain motor neuron cell bodies within the spinal cord.

## Introduction

During embryonic development, spinal motor neurons are clustered in nuclei in the ventral spinal cord, where they are segregated into distinct motor pools (Price et al., 2002). These motor neuron cell bodies project their axons out through the basement membrane boundary to innervate their peripheral targets. Many lines of evidence suggest that correct neuronal migration is required for proper settlement of cell bodies along the spinal cord to form their functional circuits (Feldner et al., 2005; Lee and Song, 2013). However, little is known about the molecular mechanisms that retain motor neuron cell bodies inside the ventral spinal cord, yet selectively allows axons to project through the basement membrane toward their peripheral targets.

The laminin-containing basement membrane plays a role not just as a barrier but also as a substrate for neuronal migration. Indeed, facial branchial motor (FBM) neurons emigrate out through holes in the basement membrane of zebrafish laminin mutants (Grant and Moens, 2010), showing that defects in the basement membrane lead to ventral mis-migration. In addition, mouse mutations for tissue-specific deletion of integrin-linked kinase or focal adhesion kinase disrupt cell basement membrane interactions or basement membrane integrity and thus these mutations cause abnormal neuronal migration (Beggs et al., 2003; Niewmierzycka et al., 2005). Together, these studies suggest that the basement membrane integrity is one of the key determinants for restricting neurons from ectopic migration.

Dystroglycan (DG) is one class of major basement membrane proteins (Bello et al., 2015). Dystroglycans are transmembrane glycoproteins, which form links between the extracellular matrix and the cytoskeleton. Glycosylated DG is necessary for ligand binding including laminin and dystrophin. Extracellular matrices directly bind to the extracellular protein, α-DG (Bello et al., 2015). This interaction results in the binding of the intracellular domain of β-DG to cytoskeleton networks or signaling components. β-DG directly binds to dystrophin (Bello et al., 2015), which is absent in Duchenne muscular dystrophy. DG is also required for precise axon guidance cue distribution and function in the nervous system (Lindenmaier et al., 2019; Wright et al., 2012). Indeed, mutation of DG showed abnormal formation of the descending hindbrain axonal tracts and defasciculation of the spinal cord dorsal funiculus (Wright et al., 2012). Furthermore, post-crossing trajectories of commissural axons were disrupted in DG mutants, similar to errors previously shown in Slit or Robo mutants. Interestingly, glycosylated DG was necessary for Slit-mediated axon guidance by regulating Slit distribution in the floor plate and the basement membrane (Wright et al., 2012). Together, these studies imply that DG is essential for correct axon guidance in the nervous system.

Several lines of evidence showed that a unique subset of neural crest cells, boundary cap (BC) cells, also control motor neuron migration. These cells migrate to the ventral motor exit points and dorsal sensory entry points and act as a barrier to motor neuron cell bodies while selectively letting their axons pass through the pial surface to the periphery. The ablation of BC cells leads to emigrant motor neurons outside the neural tube at the ventral exit points, suggesting these neural crest cells are necessary for keeping motor neurons inside the neural tube. Repulsive Semaphorin signals from BC cells prevent motor neuron cell bodies to translocate into the peripheral nervous system (PNS) (Bron et al., 2007; Mauti et al., 2007). Other signals produced by BC cells include a new Netrin family member, Netrin5, and Netrin5/DCC signals also prevent motor neuron migration out of the neural tube (Garrett et al., 2016). We previously showed that the major floor plate repellent signal, Slit2, which is also expressed in motor neurons, was downregulated in motor neurons in Islet mutants, and that Slit2 mutations cause mis-position of motor neurons in the spinal cord (Lee et al., 2015). In addition, when Slit/Robo signals were absent, motor neurons migrated out throughout the basement membrane in various levels of the spinal cord. Interestingly, in Robo1/2 double mutants, motor neurons migrated out the spinal cord at embryonic day 9.75 (E9.75) when the BC cells have not yet appeared at the ventral exit points, implying that BC cell-independent signals, including Slit/Robo signals, are required for retaining motor neurons inside the neural tube (Lee et al., 2015).

In the current study, we have therefore investigated how Slit/Robo signals are involved in preventing emigrant motor neurons in the spinal cord, by using a variety of mouse genetic tools. Our results suggest that when Slit/Robo signals were absent, especially when the signals were removed from motor neurons, that motor neuron cell bodies migrated out of the spinal cord. Moreover, this ectopic migration was associated with disruption of basement membrane and abnormal neuroepithelial organization. Together, this project reveals novel mechanistic insights of Slit/Robo signals to regulate the basement membrane integrity which is essential for restricting emigration of spinal motor neurons.

## Materials and Methods

### Mouse embryos

Mouse experiments were carried out in accordance with the National Institutes of Health Guide for the Care and Use of Laboratory Animals, by protocols approved by the University of Nevada, Reno Institutional Animal Care and Use Committee. Embryonic day 9.5 (E9.5), 10.5 (E10.5), and 12.5 (E12.5) embryos were obtained via uterine dissection of timed pregnancies, with E0.5 specified as the morning when a vaginal plug was observed. Wildtype CD-1 mice (6-8 weeks old) were purchased from Charles River Laboratories (Wilmington, MA USA). The Robo (Grieshammer et al., 2004; Long et al., 2004; López-Bendito et al., 2007), and Slit (Plump et al., 2002) mutant strains were gifts of Marc Tessier-Lavigne, Genentech and Stanford. Robo and Slit PCR genotyping were performed as previously described (Grieshammer et al., 2004; Long et al., 2004; López-Bendito et al., 2007; Plump et al., 2002). The Islet-1^MN^-GFP-F strain was a gift of Samuel Pfaff (Lewcock et al., 2007), Salk Institute, and was crossed into CD1, Robo1/2 mutant backgrounds. The Islet-Cre strain was a gift of Thomas Gould, University of Nevada, Reno. Slit2^flox^ (Rama et al., 2015) and Robo1;2^flox^ (Lu et al., 2007) mice were a gift of Le Ma, Thomas Jefferson University and Alain Chedotal, INSERM.

### Immunohistochemistry

For cryostat section immunolabeling, embryos were embedded in 15% sucrose /7.5% gelatin solutions, frozen, and then sectioned at 16 μm for E9.5 through E12.5 spinal cords using a cryostat (Leica). To melt gelatin off of tissue sections, slides were placed in warm (37-45°C) 0.1 M phosphate buffer for a couple of minutes. Sections were washed for 30 min to an hour in PBS containing 1% normal goat serum and 0.1% Triton X-100 (PBST). Primary antibodies [rabbit anti-βIII-tubulin (Covance. 1:1000), goat anti-Robo2 (R & D, 1:200), mouse anti-Islet-1 (DSHB, 1:100), mouse anti-Nestin (DSHB, 1:100), mouse anti-α-Dystroglycan (DSHB, 1:100), mouse anti-β-Dystroglycan (DSHB, 1:100)] were applied in PBST, and then slides incubated in a humidified chamber for 4 hours to overnight. After washing several hours in PBST, secondary antibodies (Jackson Immuno Laboratories) were applied for 2 hours, followed by several washes.

### In situ hybridization

Slit2 deletion in motor neurons was validated by *in situ* hybridization with a *Slit2* exon 8 - specific riboprobe. *In situ* hybridization was carried out using standard procedures (Mastick et al., 1997). The probe for Slit2 exon 8 was provided by Alain Chedotal (Rama et al., 2015), Sorbonne University, Paris, France.

### Quantification of ectopic motor neuron defects

The number of ectopic motor neurons was measured at the brachial level of the spinal cord to consistently compare an equivalent level of the spinal cord. The measurements were made on βIII-tubulin and Islet-1 labeled sections.

To determine fasciculation of exit points in the spinal cord, the width of exit points was measured at the location where motor axons exit at the brachial level of the E10.5 spinal cords. The measurements were made on βIII-tubulin labeled sections. TIFF images were imported into Image J to measure the width of motor axon exits.

The intensity of α and β-DG expression in the spinal cord was measured both in the ventral spinal cord and the dorsal spinal cord. The ventral part was determined from the ventral midline to the dorsal border of the motor nucleus, and the dorsal part was defined from the motor nucleus up to the roof plate. The measurements were made on α and β-DG labeled sections. TIFF images were imported into Image J and freehand lines were applied along the basement membrane to measure the intensity.

To measure the arrangement of neuroepithelial/radial glial processes in the ventral motor column, the distance and angle between Nestin+ fibers were measured. TIFF images of Nestin-labeled sections at E10.5 were imported and the distance and angle were measured using Image J. The measurements for each angle were presented in Rose histograms using Matlab Version 8.5.0.197613 (R2015a). Angles of each Nestin+ fibers were clustered in 10° bins. The length of each segment represented the percentage of total angles per bin.

All image analysis was conducted by an observer blind to the genotype. Data are expressed as means ± S.E.M, and differences tested for significance using student *t*-tests to analyze two groups. Data sets were tested for significance using ANOVA with Tukey’s post hoc tests to analyze multiple groups. Data are considered significantly different from the control values were when *P* < 0.05.

## Results

### Motor neurons migrate outside the spinal cord when Slit/Robo signals are missing

Spinal motor neurons cluster in clearly defined motor columns within the ventral neural tube, then project axons to innervate their peripheral targets. The main goal of our study was to examine how Slit/Robo signals keep motor neuron cell bodies within the neural tube.

Robo1 and Robo2 were expressed by spinal motor neurons and Slit2 were also expressed by the floor plate and spinal motor neurons (Brose et al., 1999; Kim et al., 2017b; Lee et al., 2015). These expression patterns of Slits and Robos suggest that motor neurons have Robo receptors to respond to Slits, which in turn could be derived from motor neurons themselves, or from the floor plate.

To observe how motor neurons are retained inside the neural tube more directly, we used the motor neuron-specific transgenic reporter Isl1^MN^-GFP (Lewcock et al., 2007) in a Robo1/2 mutant background. In wild type embryos, on E9.5, at the earliest stage of motor axons exiting the spinal cord, the Isl1-GFP+ marker clearly showed that motor neuron cell bodies were located inside the ventral spinal cord (Fig.1A). Also, no Isl1-GFP^+^ motor neuron cell bodies or axonal processes were found in the floor plate of wild type embryos (Fig. 1A). However, in *Robo1^-/-^;2^-/-^* embryos, the Isl1-GFP^+^ marker clearly showed that motor neuron cell bodies and their axonal processes were in the floor plate of the spinal cord. Also, we found that GFP^+^ motor neuron cell bodies emigrated out of the neural tube (Fig. 1B). Similarly, βIII-tubulin and Islet1 labeling on E10.5 spinal cord sections clearly showed that in wild type embryos, Islet1^+^ motor neurons were positioned the ventral spinal cord next to the floor plate and these cells were located inside the neural tube (Fig. 1C, E, and G). In contrast, we found that motor neurons streamed out of the spinal cords of E10.5 *Robo1^-/-^;2^-/-^* (Fig. 1D, F, and G). The number of ectopic motor neurons were significantly increased in *Robo1^-/-^;2^-/-^* compared to wild type or *Robo1^+/-^;2^+/-^* spinal cords, showing that Robo function is critical to prevent emigrant motor neurons (Fig. 1I). Also, the thickness at motor exit showed that *Robo1^-/-^;2^-/-^* had expanded exit points compared to wild-type or *R1^+/-^;2^+/-^* spinal cords (Fig. 1H), indicating that the motor axons exited not just in a normal narrow area but over a broader area.

**Figure 1.**
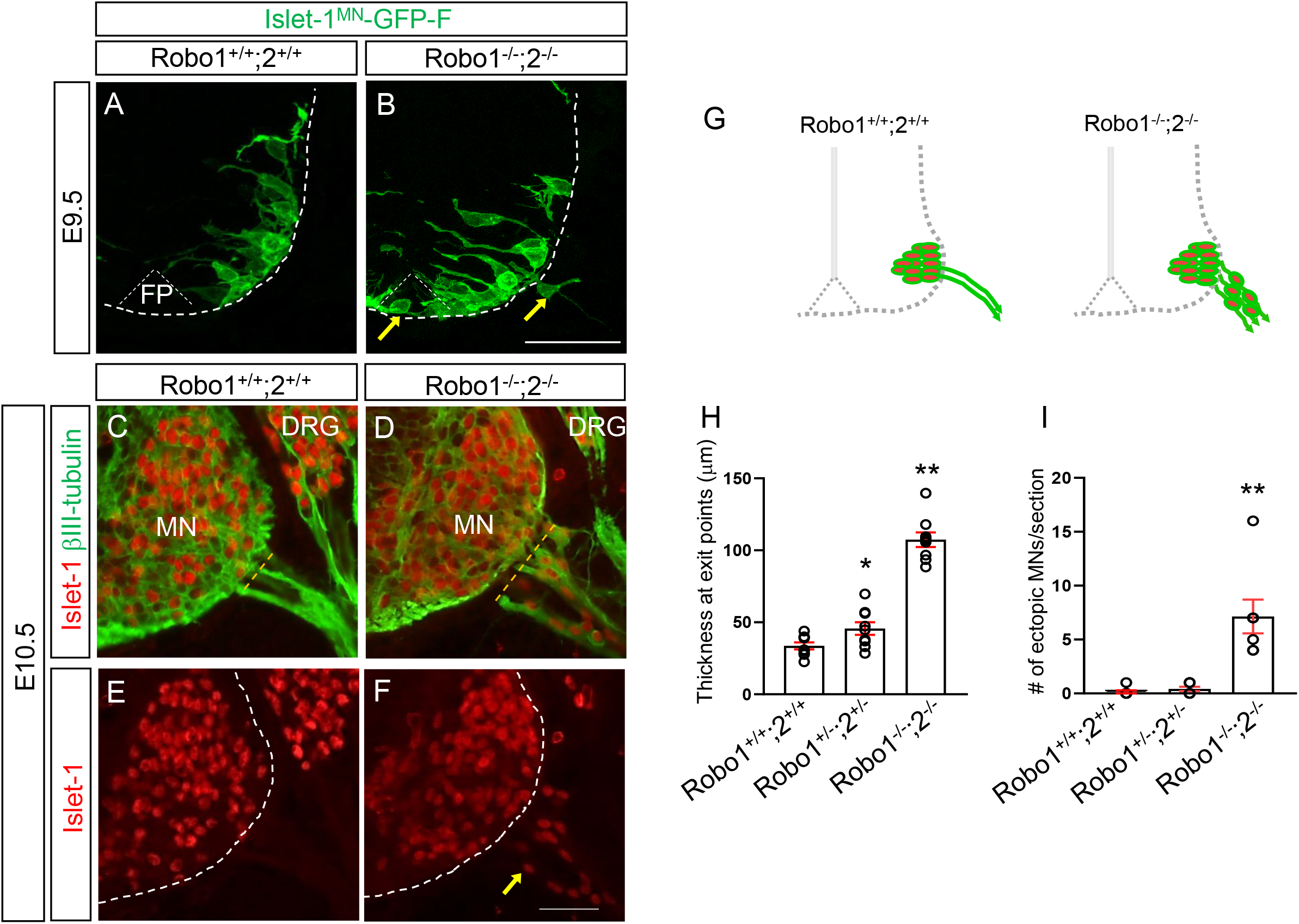
Islet-1^+^ Motor neurons migrate outside the spinal cord in Robo1/2 mutants. (A, B) Spinal cord sections of *Robo1^+/+^;2^+/+^::Islet-1^MN^-GFP-F* and *Robor1^-/-^;2^-/-^::Islet-1^MN^-GFP-F* E9.5 embryos (n=4 embryos for each genotype) show that GFP+ motor neurons are positioned within their motor column in *Robo1^+/+^;2^+/+^::Islet-1^MN^-GFP-F* (A). Arrows in B show that GFP+ motor neurons migrate into the floor plate (FP) and emigrate out of the *Robo1^-/-^;2^-/-^::Islet-1^MN^-GFP-F* spinal cord. (C-F) Islet-1 and βIII-tubulin labeling (C, D) or Islet-1 labeling (E, F) on E10.5 spinal cord sections of Robo1^+/+^;2^+/+^ and *Robo1^-/-^;2^-/-^* embryos showing that Islet-1+ motor neurons emigrate outside the *Robo1^-/-^;2^-/-^* spinal cord. (G) Schematics showing how motor neurons positioned in the E10.5 *Robo1^+/+^;2^+/+^* and *Robo1^-/-^;2^-/-^* spinal cords. (H, I) Summary graphs show the thickness at motor exit points. The width of exit points was measured at the brachial level of the E10.5 spinal cords (dotted yellow lines in C and D) (H) and number of emigrating motor neurons (I) in *Robo1^+/+^;2^+/+^, Robo1^+/-^;2^+/-^* or *Robo1^-/-^;2^-/-^* embryos (n=7 embryos for each genotype). Both thickness and ectopic motor neurons are significantly increased in Robo mutants compared to their littermate controls. Scale bars: A, B, 50 μm. C-F, 50 μm. FP, floor plate, MN, motor neuron, DRG, dorsal root ganglion. *: *P* <0.05, **: *P* <0.001.

To investigate which Robo receptors are necessary for this positioning mechanism, we used the single mutants, *Robo1^-/-^* or *Robo2^-/-^*. In *Robo1^-/-^*, βIIItubulin and Islet1 labeling on E10.5 spinal cord sections clearly showed that the number of ectopic motor neurons was not significantly different from *Robo1^+/-^* embryos (Fig. 2A-D). However, in *Robo2^-/-^*, the number of ectopic motor neurons was significantly increased compared to *Robo2^+/-^* embryos (Fig. 2E-H), suggesting that Robo2 is important for retaining motor neurons inside the spinal cord. In contrast, the thickness at the exit points were increased in *Robo1^+/-^* compared to *Robo2^+/-^* (Fig. 2R), showing that Robo1 is more important for forming fasciculated exit points in the spinal cord.

**Figure 2.**
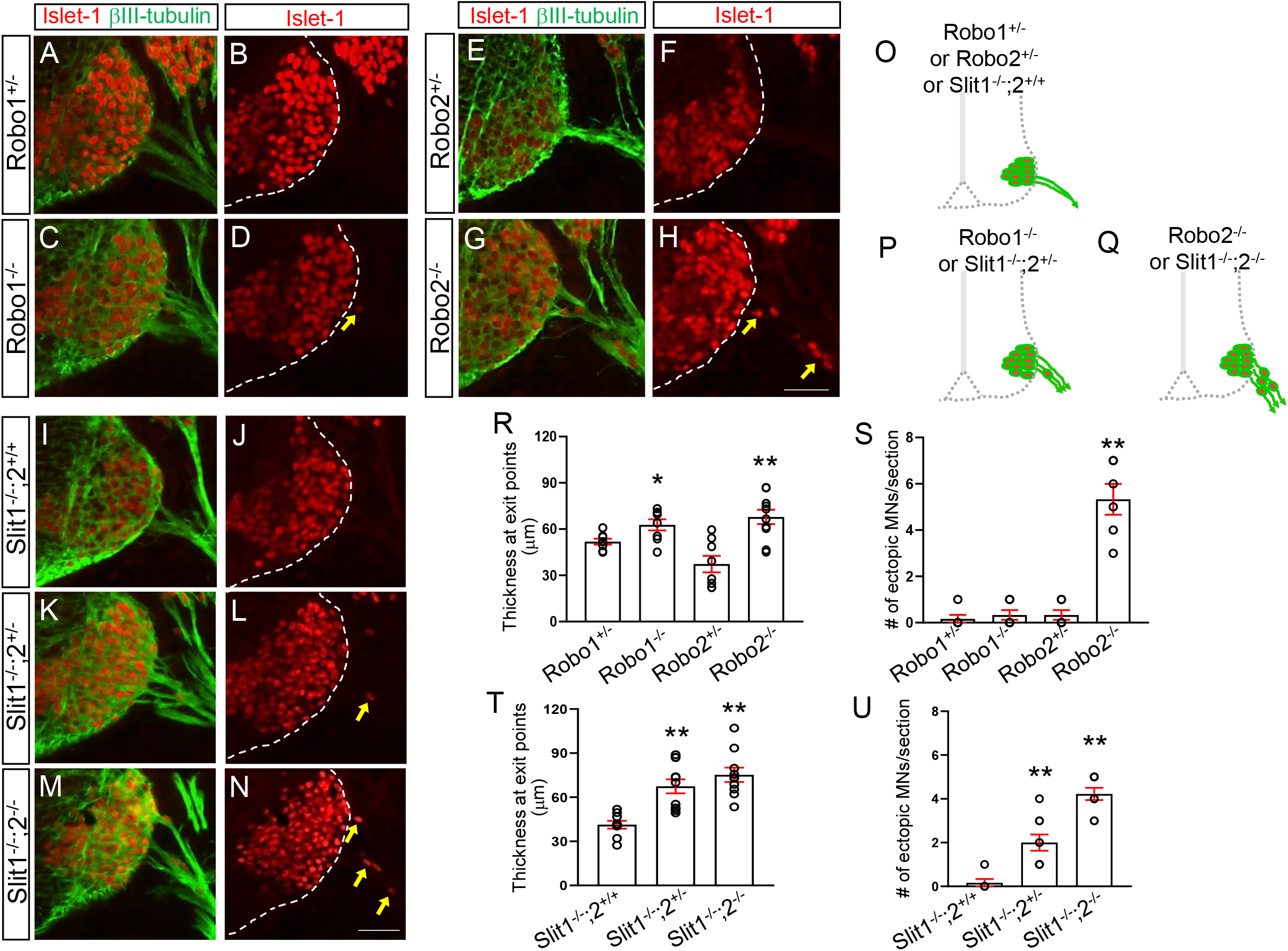
Motor neurons migrate outside the spinal cord in Robo2 mutants and Slit1/2 mutants. (A-H) Islet-1 and βIII-tubulin labeling (A, C, E, G) or Islet-1 labeling (B, D, F, H) on E10.5 spinal cord sections of *Robo1^+/-^, Robo1^-/-^, Robo2^+/-^*, and *Robo2^-/-^* (n=7 embryos for each genotype) showing that significant numbers of Islet-1+ motor neurons emigrate outside the *Robo2^-/-^* spinal cord. (I-N) Islet-1 and βIII-tubulin labeling (I, K, M) or Islet-1 labeling (J, L, N) on E10.5 spinal cord sections of *Slit1^-/-^;2^+/+^, Slit1^-/-^;2^+/-^* or *Slit1^-/-^;2^-/-^* (n=8 embryos for each genotype) showing that significant numbers of Islet-1+ motor neurons emigrate outside the *Slit1^-/-^;2^-/-^* spinal cord. (O-Q) Schematics showing how motor neurons positioned in the E10.5 *Robo1^+/-^, Robo1^-/-^, Robo2^+/-^, Robo2^-/-^ Slit1^-/-^;2^+/+^*, *Slit1^-/-^;2^+/-^* or *Slit1^-/-^;2^-/-^* spinal cords. (R, S) Summary graphs show the thickness at motor exit points (R) and number of emigrating motor neurons (S) in single Robo mutants and their littermate controls. In *Robo1^+/-^* and *Robo1^-/-^* embryos, Islet-1+ motor neurons are within the spinal cord even though these embryos have wider motor exits. However, both thickness and ectopic motor neurons are significantly increased in *Robo2^-/-^* compared to their littermate controls. (T, U) Summary graphs show the thickness at motor exit points (T) and number of emigrating motor neurons (U) in Slit1/2 mutants and their littermate controls. *Slit1^-/-^;2^+/-^* or *Slit1^-/-^;2^-/-^* embryos have extended motor exits compared to *Slit1^-/-^;2^+/+^* embryos. The number of ectopic motor neurons are significantly increased in *Slit1^-/-^;2^-/-^* than *Slit1^-/-^;2^+/-^ or Slit1^-/-^;2^+/+^* embryos. Scale bars: A-H, 50 μm, I-N, 50 μm. *: *P* <0.05, **: *P* <0.001.

We also tested whether Slit mutant embryos have the same phenotypes observed in Robo mutant embryos. We found that spinal motor neurons streamed out the neural tube when both Slit1 and Slit2 were missing (Fig. 2M, N). Also, in Slit1/2 double mutants, the thickness of the exit points was significantly increased compared to *Slit1^-/-^;2^+/+^* (Fig. 2T). To test which Slit is important for keeping motor neurons inside the spinal cord, Slit single mutants, *Slit1^-/-^* or *Slit2^-/-^*, were used. In *Slit1^-/-^;2^+/+^* mutants, no ectopic motor neurons were found outside of the spinal cord (Fig. 2I, J and U). In contrast, emigrant spinal motor neurons were found in *Slit2^-/-^* embryos (Fig. 3L, M and V), suggesting that Slit2 is critical for retaining motor neurons inside the neural tube. Interestingly, extended exit points were also observed when a single wildtype *Slit2* allele was present in *Slit1^-/-^;2^+/-^* (Fig. 2K, L). However, we still found one or two emigrant motor neurons in the *Slit1^-/-^;2^+/-^* mutants even though one copy of Slit2 prevented the significant number of motor neurons streaming out the neural tube observed in Slit1/2 double mutants. Overall, our main finding is that Slit2 is required for keeping motor neurons within the spinal cord.

**Figure 3.**
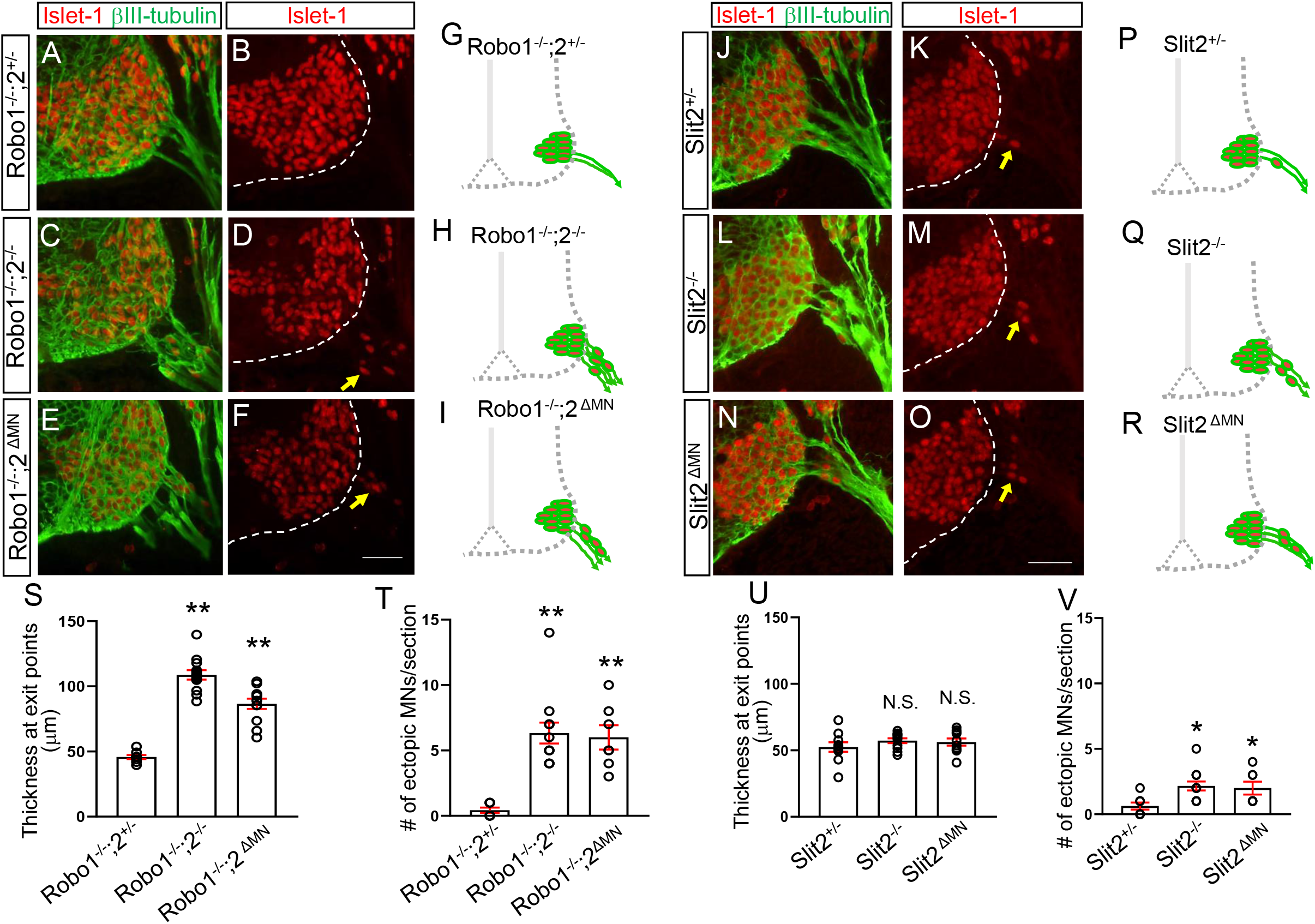
Motor neuron-specific Robo2 or Slit2 knockout mutants lead to emigrant motor neurons in the spinal cord. (A-F) Islet-1 and βIII-tubulin labeling (A, C, E) or Islet-1 labeling (B, D, F) on E10.5 spinal cord sections of *Robo1^-/-^;2^+/-^, Robo1^/-^;2^-/-^* or *Robo1^-/-^;2^ΔMN^* (n=8 embryos for each genotype) showing that significant numbers of Islet-1+ motor neurons emigrate outside the *Robo1^-/-^;2^-/-^* or *Robo1^-/-^;2^ΔMN^* spinal cord. (G-I) Schematics showing how motor neurons positioned in the E10.5 *Robo1^-/-^;2^+/-^, Robo1^-/-^;2^-/-^* or *Robo1^-/-^;2^ΔMN^* embryos. (J-O) Islet-1 and βIII-tubulin labeling (J, L, N) or Islet-1 labeling (K, M, O) on E10.5 spinal cord sections of *Slit2^+/-^, Slit2^-/-^* or *Slit2^ΔMN^* (n=8 embryos for each genotype) showing that significant numbers of Islet-1+ motor neurons emigrate outside the *Slit2^-/-^ or Slit2^ΔMN^* spinal cord. (P-R) Schematics showing how motor neurons positioned in the E10.5 *Slit2^+/-^, Slit2^-/-^* or *Slit2^ΔMN^* embryos. (S. T) Summary graphs show the thickness at motor exit points (S) and number of ectopic motor neurons (T) in *Robo1^-/-^;2^+/-^*, *Robo1^-/-^;2^-/-^* or *Robo1^-/-^;2^ΔMN^* embryos. Both motor exit points and the number of ectopic motor neurons are significantly increased in *Robo1^-/-^;2^ΔMN^* embryos compared to *Robo1^-/-^;2^+/-^* embryos. (U, V) Summary graphs show the thickness at motor exit points (U) and number of ectopic motor neurons (V) in *Slit2^+/-^, Slit2^-/-^* or *Slit2^ΔMN^*embryos. The number of ectopic motor neurons is significantly increased in *Slit2 ^ΔMN^* embryos compared to *Slit2^+/-^* embryos. Scale bars: A-F, 50 μm, J-O, 50 μm. *: *P* <0.05, **: *P* <0.001.

### Motor neuron-specific Robo2 or Slit2 knockout mutants lead to emigrant motor neurons in the spinal cord

The emigrant phenotype might be due to indirect effects of global knockout of Robo receptors. A stringent genetic test for Robo-dependent function of motor neurons in positioning mechanism would be to use motor neuron-specific knockouts. To determine the cell autonomy of Slit reception in these cells, the Isl1-cre line was used to delete Robo signaling in motor neurons in *Robo1^-/-^; Robo2^flox/-^* embryos. *Robo1^-/-^; Robo2^flox/-^* was crossed to *Robo1^+/-^; Robo2^+/-^; Tg Isl1-Cre*, to produce Robo2^ΔMN^ embryos. Robo2 antibody labeling was used to confirm Robo2 loss in motor neurons (Supplementary Fig. 1). The Robo2^ΔMN^ spinal cords have substantial motor neuron emigration, in which the number of ectopic motor neurons was not significantly different from Robo1/2 global knockout embryos (Fig. 3E, F, I and T). In contrast, no emigrant motor neurons were found in the *Robo1^-/-^;2^+/-^* spinal cords (Fig. 3A, B, G, and T). Moreover, the Robo2^AMN^ spinal cords had defasciculated exit points which were not observed in the *Robo1^-/-^;2^+/-^* spinal cords (Fig. 3E, S). These results suggest that motor neurons are the essential site for Robo function to keep motor neurons inside the spinal cord.

Since Slit2 is expressed by motor neurons, we did a direct genetic test by motor neuron-specific knockout of Slit2. The Isl1-cre line was used to delete Slit2 in *Slit2^flox/-^* embryos. *Slit2^flox/-^* was crossed to *Slit2^+/-^; Tg Isl1-Cre* to get the desired Slit2^ΔMN^ embryos. These embryos were confirmed by Slit2 *in situ* hybridization to have a motor neuron-specific reduction of Slit2 exon8 transcripts, while retaining wild type levels of Slit2 transcripts in the floor plate, although the Slit2^-^ allele does produce a low level of exon 8-containing transcripts (Supplementary Fig. 2). The Slit2^ΔMN^ spinal cords had motor neuron emigration, in which the number of ectopic motor neurons was not significantly different from Slit2 global knockout embryos (Fig. 3N, O, R and V). This implies that motor neurons are the critical site for Slit/Robo signaling and Slit2 produced by motor neurons is important for preventing motor neuron emigration. In contrast, the number of emigrant motor neurons was significantly less in the *Slit2^+/-^* spinal cords (Fig. 3J, K, P, and V). However, we found wider exit points in the *Slit2^+/-^* spinal cords as well as the Slit2^ΔMN^ spinal cords (Fig. 3U), showing that Slit2 is important for regulating fasciculated exits in the spinal cord.

Together, these data show that cell autonomous signaling by Slit/Robo in spinal motor neurons play a significant role in keeping motor neurons inside the spinal cord.

### Dystroglycan expression is discontinuous and ruptured in Robo or Slit mutants

Because motor neurons are able to emigrate through the basement membrane in Slit and Robo mutants, we examined the basement membrane integrity in the spinal cord. We first examined patterns of Dystroglycan (DG), the major basement membrane proteins expressed in the spinal cord.

Firstly, the intensity of α-DG expression was measured after immunostaining on spinal cord sections. In Robo1/2 double knockout spinal cords, we found that the basement membrane was discontinuous, specifically in the ventral spinal cord (Fig. 4B, E). Strikingly, gaps in the basement membrane were associated with the ventral sites where motor neurons ectopically emigrate, suggesting that emigrating motor neurons were associated with degraded basement membrane. Similarly, β-DG expression was also significantly decreased in the knockout spinal cords (Fig. 4N). Importantly, both α and β-DG expression remained intact in the dorsal part of the knockout spinal cord (Fig. 4M, N), consistent with the ventral Slit2 signals derived from motor neurons. Also, in *Slit2^-/-^* spinal cords, both α and β-DG antibody labeling were decreased compared to *Slit2^+/-^* (Fig. 4H, K). However, the antibody labeling was not changed in the dorsal part of the *Slit2^-/-^* spinal cord (Fig. 4O, P). This implicates that Slit proteins produced by ventral tissues are important for regulating DG protein levels in the spinal cord.

**Figure 4.**
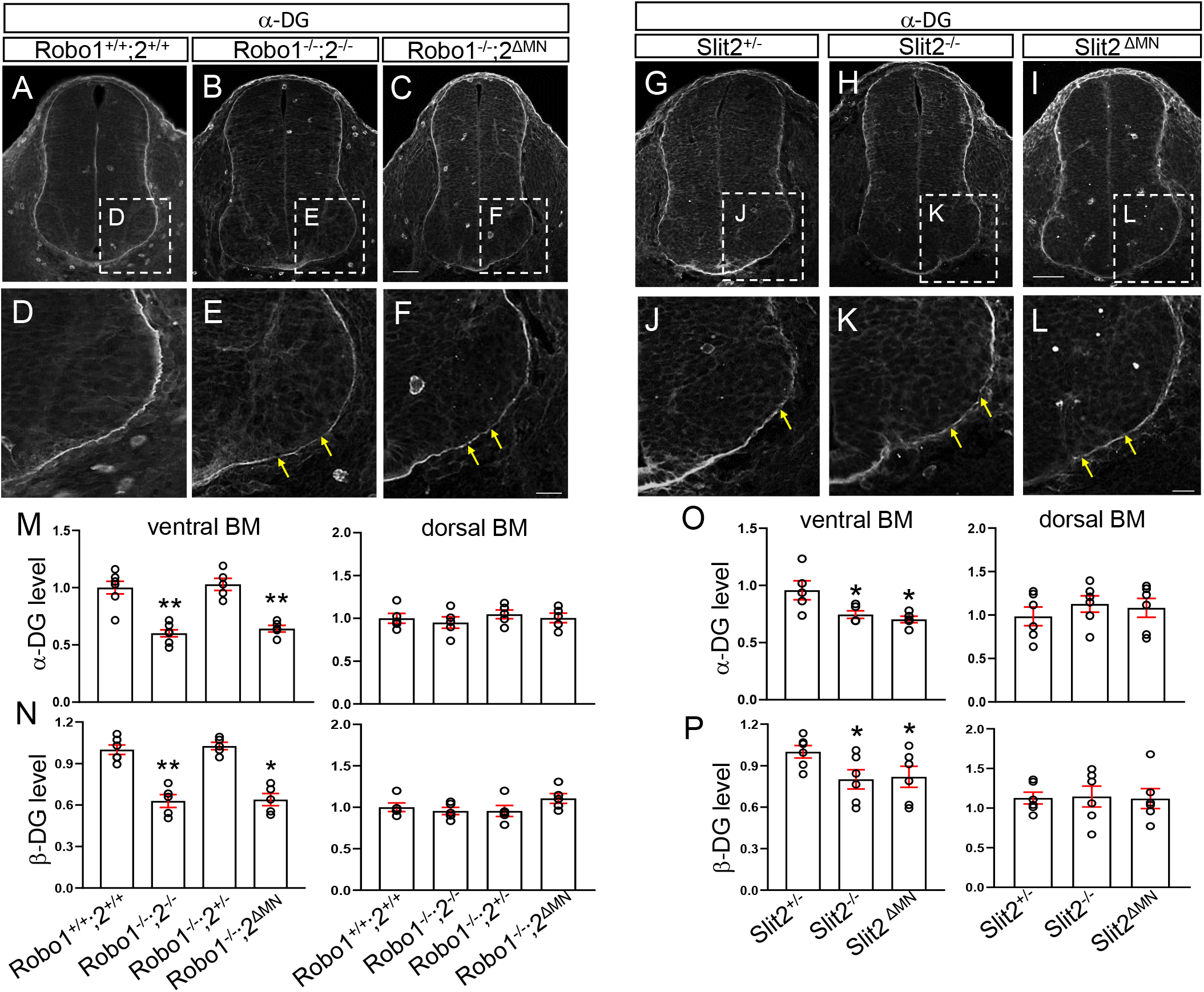
Dystroglycan ventral expression is lower and ruptured in Robo or Slit knockout spinal cords. (A-F) α-DG labeling on E10.5 spinal cord sections of *Robo1^+/+^;2^+/+^, Robo1^-/-^;2^-/-^* or *Robo1^/-^;2^ΔMN^* (n=5 embryos for each genotype). (D-F) Close-up images showing that α-DG expression is discontinuous in the ventral spinal cord of *Robo1^-/-^;2^-/-^* or *Robo1^-/-^;2^ΔMN^* embryos (yellow arrows). (G-L) α-DG labeling on E10.5 spinal cord sections in Slit2 embryos. (J-L) Close-up images showing that α-DG expression is discontinuous in the ventral spinal cord of *Slit2^-/-^* or *Slit2^ΔMN^* embryos (n=6 embryos for each genotype, yellow arrows). (M, N) Summary graphs show intensity (pixel gray level) of α-DG expression (M) and β-DG expression (N) in the ventral and dorsal part of the spinal cord. Both a and β-DG expression is significantly decreased in the ventral part of the spinal cord of *Robo1^-/-^;2^-/-^* or *Robo1^-/-^;2^A1M1N^* embryos. However, α and β-DG expression is maintained in the dorsal part of the spinal cord of *Robo1^-/-^;2^-/-^* or *Robo1^-/-^;2^ΔMN^* embryos. (O, P) Summary graphs show intensity (pixel gray level) of α-DG expression (O) and β-DG expression (P) in the ventral and dorsal part of the spinal cord. Both α and β-DG expression is significantly decreased in the ventral part of the spinal cord of *Slit2^-/-^* or *Slit2^ΔMN^* embryos. However, both α and β-DG expression is not changed in the dorsal part of the spinal cord of *Slit2^-/-^* or *Slit2^ΔMN^* embryos. Scale bars: A-C, 50 μm; D-F, 20 μm, G-I, 50 μm, J-L, 20 μm. *: *P* <0.05, **: *P* <0.001.

To further localize the site of Robo and Slit function in maintaining intact ventral basement membranes, we examined basement membrane integrity in Robo2 and Slit2 motor neuron specific knockouts, both of which cause motor neuron emigration. We first examined the DG antibody labeling patterns in Robo2^ΔMN^ spinal cord. In the mutant, both α and β-DG expression were significantly decreased compared to *Robo1^-/-^;2^+/-^* (Fig. 4C, F, M, and N). Then, we investigated Slit2^ΔMN^ embryos. Similarly, in the mutant spinal cord, DG expression in the ventral basement membrane was discontinuous. Also, both α and β-DG expression were significantly decreased, similar to *Slit2^-/-^* spinal cords (Fig. 4I, L, O, and P). However, density of the dorsal part of the basement membrane was not significantly changed in either conditional knockout spinal cords, consistent with the ventral loss of Slit2 (Fig. 4M, N, O, and P). These results suggest that DG levels and basement membrane integrity depend on Robo2 and Slit2 function in motor neurons, rather than on other ventral tissues such as the floor plate.

### Neuroepithelial endfeet organization is disrupted in Robo or Slit mutant spinal cords

Dystroglyan proteins are important for the organization of the neuroepithelial fibers which extend endfeet radially out to form the pial surface of the neural tube, in that disruption of dystroglycan levels or assembly can disrupt organization of radial fibers and endfeet (Schroder et al., 2007). Therefore, we examined the neuroepithelial radial processes and endfeet using Nestin antibody labeling on spinal cord sections of *Robo1^-/-^;2^-/-^* or motor neuron-specific Robo2 knockout embryos. In wildtype spinal cord, Nestin antibody labeling showed a radial organization of aligned fibers reaching out to the pial surface of the spinal cord. The fibers were relatively evenly spaced and had consistent angles (Fig 5A, A’ and H). When Robo1 and 2 were missing, radial processes were sparse, disorganized, and directed at a wider range of angles (Fig. 5C, C’ and H). In addition, fewer fibers reached to the pial surface. In the motor neuron-specific Robo2 knockout spinal cord, the distance between the processes was less variable than observed in the global knockout spinal cord (Fig. 5D, H). However, the processes in the mutant had high variance angles in the motor column (Fig. 5D, D’). Similar defects were seen in Slit2-/- and Slit2^ΔMN^ mutant spinal cords (Fig. 5F, F’, G, G’, and I). Overall, these results add to the observations that motor neuron emigration is associated with abnormal basement membrane formation and organization of radial neuroepithelial fibers and endfeet.

**Figure 5.**
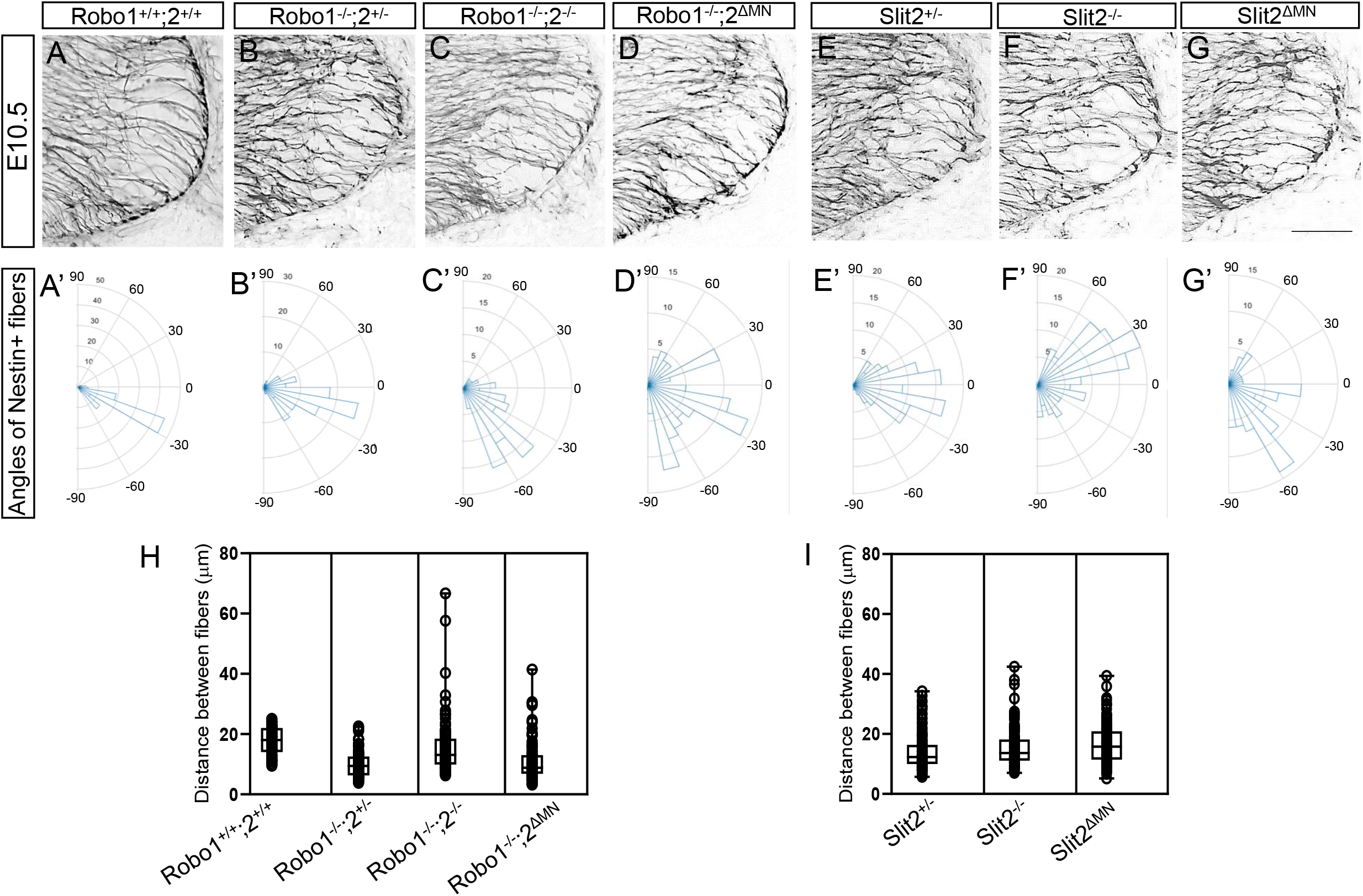
Nestin-positive neuroepithelial cells are disorganized in Robo or Slit2 knockout spinal cords. (A-G) Nestin labeling on E10.5 (n=6 embryos for each genotype) spinal cord sections in Robo (A-D) and Slit (E-G) embryos showing that Nestin+ neuroepithelial/radial glial processes were sparse, disorganized, and disrupted in the ventral spinal cord of *Robo1^-/-^;2^-/-^, Robo1^-/-^;2^ΔMN^, Slit2^-/-^ or Slit2^ΔMN^* embryos. embryos. (A’-G’) Rose histograms showing the distributions of angles inside the spinal motor column. Angles of each projection were clustered in 10° bins. The length of the radius of each segment represents the percentage of total fibers include each bin. Note that the radial lengths were shown on a shorter scale in the mutant graphs because the angles were more variable than wild-type embryos, and therefore had lower percentages per bin. (H, I) Nested box-and-whisker plots with aligned data points showed that distance between Nestin+ processes has a high variance in *Robo1^-/-^;2^-/-^* or *Robo1^-/-^;2^ΔMN^, Slit2^-/-^* or *Slit2^ΔMN^* embryos compared to wild-type or their littermate controls. Scale bars: A-F, 50 μm. *: *P* <0.05, **: *P* <0.001.

## Discussion

The basement membrane boundary between the CNS and PNS is critical for proper spinal cord development and function, as exemplified by the proper positioning of spinal motor neuron cell bodies for their functional input and output. However, little is known about how motor neurons keep their organized pattern inside the ventral spinal cord. In the current study, we investigate the molecular basis of how spinal motor neurons are retained within the neural tube by using of mouse genetic tools.

Our main finding is that Slit/Robo signals are necessary to keep spinal motor neurons inside the neural tube by regulating the basement membrane integrity. The genetic approaches with different combinations of mutants suggest that motor neurons are the essential site for Slit/Robo signaling to set up and maintain a normal basement membrane in the spinal cord.

Importantly, motor neuron-specific Robo or Slit knockout results demonstrate the cell-autonomous requirement for Slit/Robo signaling in keeping motor neuron within the neural tube. It is interesting to note that the floor plate function was not necessary to retain motor neurons in the spinal cord. Contrast to the previous reports showing a ventral repulsive role in controlling the position of the motor neurons within the neural tube, prevention attraction into the floor plate (Kim et al., 2017a; Kim et al., 2015), and in setting the position of the motor exit point (Kim et al., 2017b), emigration does not neatly fit into a push-pull balance between midline attraction and repulsion. The results instead point to a local role in controlling the ability of motor neurons to either stay in their normal place in the nucleus, or emigrate out. A local role of Slit/Robo signaling is strongly implicated by the cell autonomous functions shown by the motor neuron specific knockouts. In addition, our findings suggest that motor neurons have a strong migratory ability, and apparently can follow their axons out into the periphery, if Slit/Robo signals are disrupted.

### Basement membrane integrity is essential for proper neuronal migration

Basement membrane integrity is one of the key determinants for proper neuronal migration. Dystroglycan (DG) is the major basement membrane protein, and this protein is important for connection between the extracellular matrix and the cytoskeleton (Bello et al., 2015). In addition, DG is important for regulating neuronal migration and organizing guidance cues in their precise location (Lindenmaier et al., 2019; Wright et al., 2012). One of our key findings is that when Slit/Robo signals are absent, motor neurons migrate out of the spinal cord, and that this abnormal emigration is associated with disruptions in DG expression in the ventral neural tube. Interestingly, both α and β-DG expression remained intact in the dorsal part of the knockout spinal cord (Fig. 4). This implicates that Slit proteins produced in ventral tissues are important for regulating DG expression in the spinal cord. The results from the motor neuron-specific Robo2 and Slit2 knockout spinal cords support this finding that motor neurons located at the ventral part of the spinal cord are one of the key sites that Slit/Robo signals act to set up a normal basement membrane. It is possible that Slit2 protein produced by ventral tissues, including locally by the motor neurons themselves, accumulates in the basement membrane through interaction with DG (Wright et al., 2012). This potential accumulation of Slit2 at the basement membrane may function in a repulsive manner to prevent motor neuron emigration. Thus, motor neuron-derived Slit2 could make an additional barrier to repel motor neurons near the exit points. On the other hand, emigrant motor neurons could play an active degrading role: They are released to migrate in Robo and Slit mutants, and then the migrating motor neurons are able to degrade the basement membrane and escape (Santiago-Medina et al., 2015).

Wider motor axon exits through the basement membrane could lead to ectopic motor neurons as they penetrate through the ventral part of the basement membrane. The extracellular proteinases, such as matrix metalloproteinases (MMPs), expressed by motor axons have been implicated to play a role in axon pathfinding by degrading the extracellular matrix (McFarlane, 2003; Yong et al., 2001). Thus, a wider front of motor axons penetrating the basement membrane could provide a route for ectopic migration, possibly by degrading broad regions of the basement membrane. However, even though *Robo1^+/-^* and *Robo1^-/-^* had wider exits, the number of ectopic motor neurons was significantly reduced compared to *Robo2^-/-^*, in which the width of exit points was similar to *Robo1^+/-^* or *Robo1^-/-^*. This finding suggests that defasciculated wider motor axon exits are not sufficient to cause motor neuron emigration in the spinal cord, and simply the width of the exit point did not closely predict the severity of motor neuron emigration.

### Slit/Robo signals organize normal radial structure of the neuroepithelial cells

The neuroepithelial cells are presumably the main producers of the basement membrane as these cells have long radial glia-like extensions out to the pial surface, and their endfeet directly form the pial surface of the spinal cord. Since DG expression of the basement membrane is reduced in Slit or Robo mutants, this implies that Slit/Robo signaling must be required to stimulate the production of sufficient basement membrane components such as DG, or to maintain these basement membrane assemblies. Our results also implicate Slit/Robo signaling in organizing and aligning the neuroepithelial end feet; this Slit/Robo signaling is a quite unexpected function that may reveal a novel role in guiding these cellular projections.

DG plays a role in regulating neuroepithelial cell morphology by anchoring their endfeet to the basal lamina (Schroder et al., 2007). In the study, they reported that knockdown of DG and overexpression of a dominant-negative DG in neuroepithelial cells of chick retina lead to abnormal radial morphology of the cells (Schroder et al., 2007). Consistent with their findings, we observed that the loss of DG expression in the ventral spinal cord was associated with abnormal neuroepithelial cell morphology in global and motor neuron-specific Slit or Robo mutants. However, it remains unresolved whether Slit/Robo signals directly act on these neuroepithelial cells and affect their functions, or whether Slit/Robo signals somehow act on basement membrane assembly and then lead to an indirect effect on neuroepithelial fibers.

### Other possible mechanisms: Slit/Robo signals in maintaining the position of motor neurons within the neural tube

Semaphorin signals are also a critical factor in limiting motor cell body migration as Sema6A is expressed by boundary cap (BC) cell clusters located at motor exits. BC cells serve as a boundary between central and peripheral nervous system and Sema6A signals repel motor cell bodies that express Nrp2 or Plexins (Bron et al., 2007; Mauti et al., 2007). Knockdown or genetic deletion of Nrp2, Sema6A and PlexinA induced motor neuron emigration in a BC-cell dependent manner. Our recent study showed that Nrp1 and Slit2 was downregulated in motor neurons in Islet mutants and knockdown of both Npr1 and Slit2 caused ectopic motor neurons (Lee et al., 2015). However, little is known about how Sema and Slit signals may cooperate in maintaining the position of spinal motor neurons.

The potential interaction between Slit and Sema signaling in regulating neuronal migration and axon guidance has been reported. Indeed, *in vivo* and *in vitro* studies showed that Robo1 interacted with Nrp1 to modulate Semaphorin signaling and regulated the migration of interneurons (Andrews et al., 2006). In addition, PlexinA1 bound the C-terminal Slit fragment and transduced a SlitC signal during commissural axon guidance in the spinal cord (Delloye-Bourgeois et al., 2015). Therefore, testing the functions of Sema/Nrp signals in spinal motor neurons is an exciting path to gain insights into how Slit and Sema signals interact. Further research will be needed to define genetic interaction between Slit2 and Nrp1 by generating combined mutants and using *in vitro* assays.

Another possibility is that Slit/Robo repulsion could regulate cell adhesion during motor neuron positioning. Cadherins, calcium-dependent adhesion receptors, play an essential role in cell movement. Indeed, *in vivo* and *in vitro* study showed that N-cadherin deficient cells did not lead to cell adhesion and migration (Shih and Yamada, 2012). For example, Slit1/Robo2 signals regulate ganglion assembly through N-cadherin signals in the placodal cells (Shiau and Bronner-Fraser, 2009). A recent study showed that Slit2/Robo1 signals promoted the adhesion of tongue carcinoma cells by downregulating the expression of E-cadherin (Zhao et al., 2016). These findings suggest that Slit/Robo signals could regulate spinal motor neuron adhesion within their motor column possibly through affecting cadherin signals.

Overall, our findings show that the loss of Slit/Robo signals from motor neurons leads to ectopic motor neuron migration, and that this emigration is associated with strong disruptions of the basement membrane in the ventral spinal cord. Furthermore, our genetic approaches imply that motor neurons within the ventral spinal cord are the vital site for Slit/Robo signaling in establishing the normal basement membrane.

## Acknowledgements

The *Robo* and *Slit* mutant founder mice were gifts of Marc Tessier-Lavigne (Stanford; Genentech). The Islet-1^MN^-GFP-F mice were gifts of Samuel Pfaff (Salk Institute). The Islet-Cre strain was a gift of Thomas Gould (University of Nevada, Reno). Slit2^flox^ and Robo1;2^flox^ mice were a gift of Le Ma (Thomas Jefferson University). The probe for Slit2 exon 8 was a gift of Alain Chedotal (Sorbonne University, Paris, France). Several people in the Mastick lab provided help and discussions on this project, including Claudia Garcia-Pena, G. Eric Robinson and Sterling Louw. This project was supported by NIH RO1 EY025205 to GSM. Use of tissue culture and imaging core facilities was supported by P20 RR-016464, P20 GM103440, P20 GM103554, and P20 GM103650.

---

**Supplementary Figure 1.**
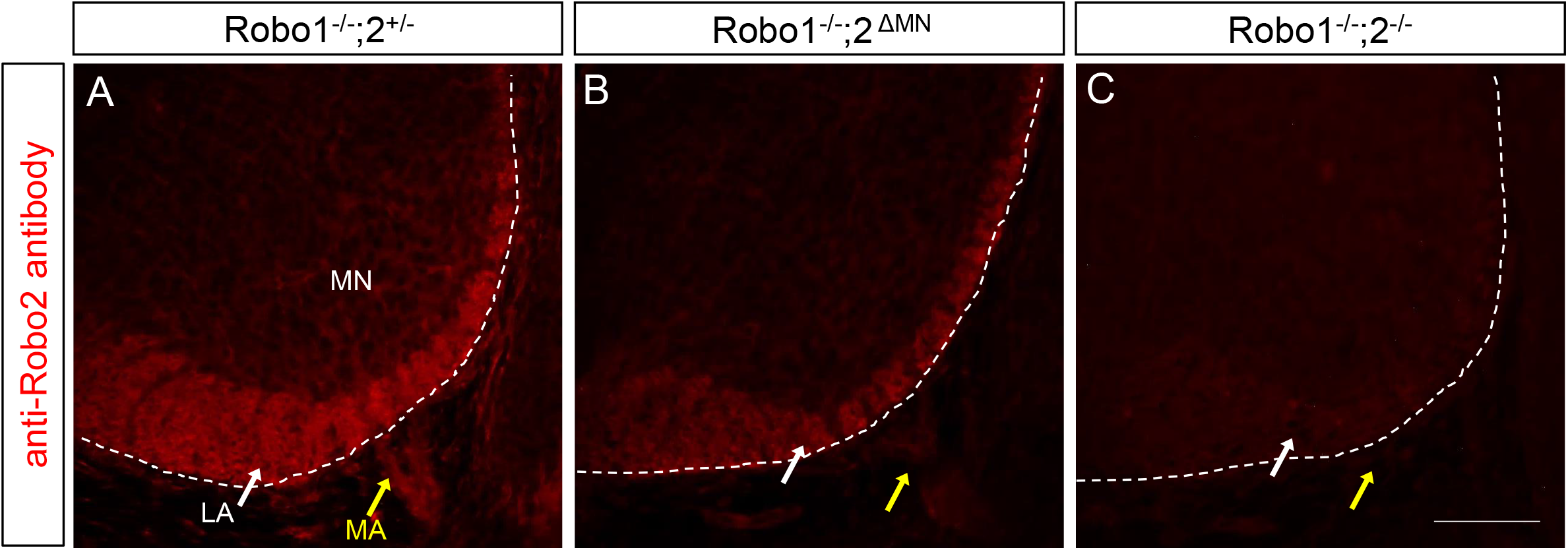
Robo2 antibody labeling is depleted by spinal motor neurons and motor axons in the motor neuron-specific Robo2 knockout spinal cord. (A-C) Robo2 labeling on E12 (n=4 embryos for each genotype) spinal cord sections in *Robo1^-/-^;2^+/-^, Robo1^-/-^;2^ΔMN^*, or *Robo1^-/-^;2*^-/-^. (A) In *Robo1^-/-^;2^+/-^*, Robo2 is expressed by lateral longitudinal axons (LA, white arrow) and motor neuron cell bodies and their axons (MA, yellow arrow). (B) In *Robo1^-/-^;2^ΔMN^*, Robo2 is only expressed by longitudinal axons and depleted by motor neuron cell bodies and their axons. (C) In *Robo1^-/-^;2^-/-^*, Robo2 expression is totally departed in the spinal cord. MN, motor neurons, LA, Longitudinal axons, MA, motor axons, Scale bars: A-C, 50 μm.

**Supplementary Figure 2.**
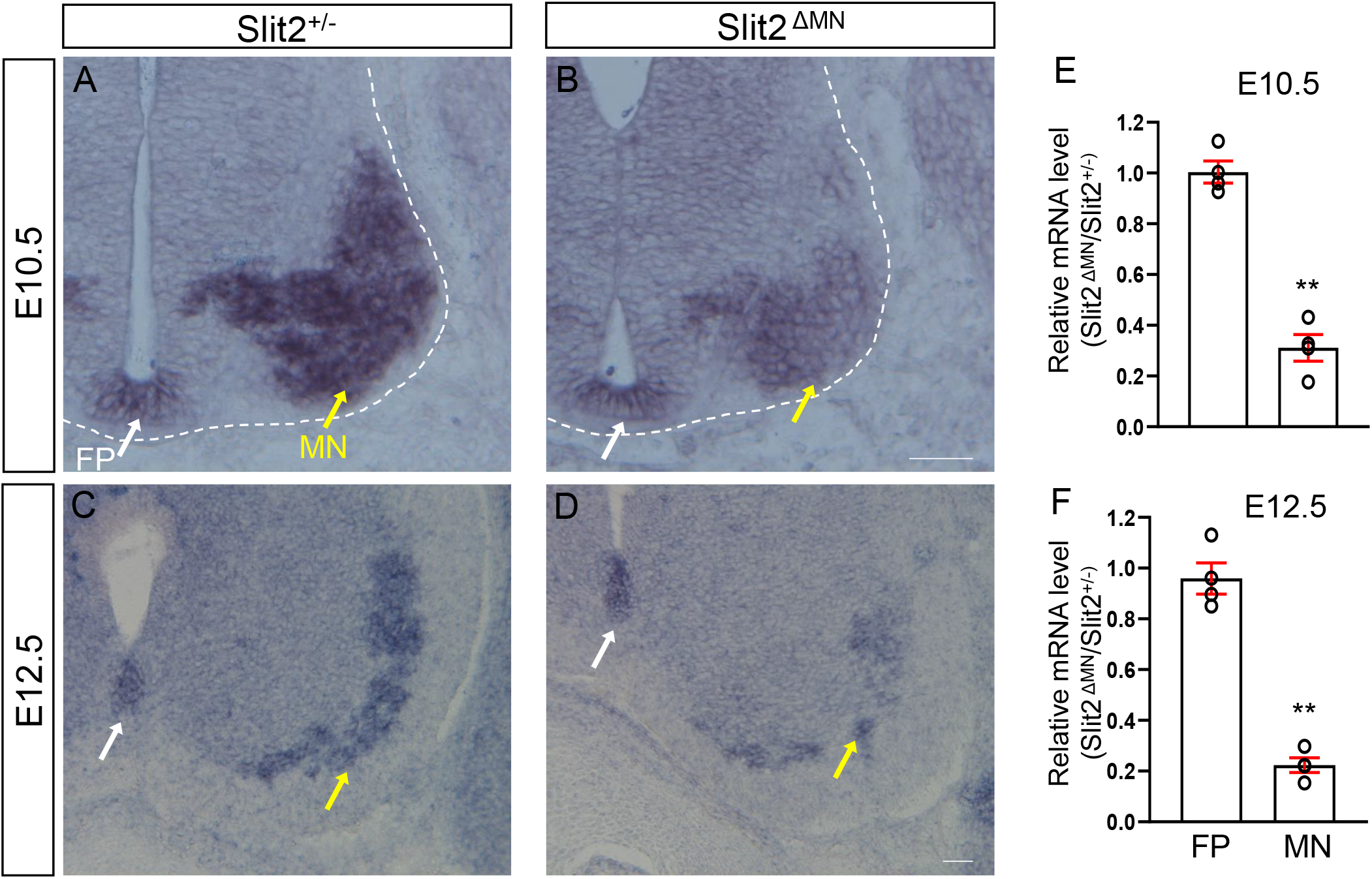
Slit2 exon 8 transcript is reduced by spinal motor neurons in the motor neuron-specific Slit2 knockout spinal cord. (A-D) In situ hybridization with Slit2 exon 8 transcript on E10.5 (A, B) and E12.5 (C, D) spinal cord sections in *Slit2^+/-^* (A, C) or *Slit2^ΔMN^*(B, D). Slit2 exon 8 transcript was labeled by the floor plate (white arrows) and motor nuclei (yellow arrows) in *Slit2^+/-^*. In *Slit2^ΔMN^*, expression of the transcript was significantly reduced by spinal motor neurons (MN) but no changes in the floor plate (FP). (E, F) Summary graphs show relative mRNA level in the floor plate (FP) and the motor neurons (MN) of E10.5 and E12.5 spinal cords (n=4 embryos for each genotype). Scale bars: A and B, 50 μm; C and D, 50 μm. **: *P* <0.001.

